# Commentary: Is the Statistic Value All We Should Care about in Neuroimaging?

**DOI:** 10.1101/064212

**Authors:** Gang Chen, Paul A. Taylor, Robert W. Cox

## Abstract

Here we address an important issue that has been embedded within the neuroimaging community for a long time: the absence of effect estimates in results reporting in the literature. The statistic value itself, as a dimensionless measure, does not provide information on the biophysical interpretation of a study, and it certainly does not represent the whole picture of a study. Unfortunately, in contrast to standard practice in most scientific fields, effect (or amplitude) estimates are usually not provided in most results reporting in the current neuroimaging publications and presentations. Possible reasons underlying this general trend include: 1) lack of general awareness, 2) software limitations, 3) inaccurate estimation of the BOLD response, and 4) poor modeling due to our relatively limited understanding of FMRI signal components. However, as we discuss here, such reporting damages the reliability and interpretability of the scientific findings themselves, and there is in fact no overwhelming reason for such a practice to persist. In order to promote meaningful interpretation, cross validation, reproducibility, meta and power analyses in neuroimaging, we strongly suggest that, as part of good scientific practice, effect estimates should be reported together with their corresponding statistic values. We provide several easily adaptable recommendations for facilitating this process.

## Introduction

Just as cartography requires a balance to be struck between the loss of important detail and the exactitude of a map that has “the scale of a mile to the mile” (Carroll, 1889), so too science rquires careful extraction and summarization following an experiment. In other words, to present concisely the important components of the data and analyses, an investigator reports the experiment and makes a generalized conclusion based on some supporting evidence: a small condensed set of numbers. The crucial question is: how much or to which extent should the investigator compress the information without sacrificing too much? There are arbitrary choices that have to be made, but there are some definite thresholds under which loss of information is too great for optimal utility.

For example, in a typical statistical analysis, two quantitative results are produced for each effect of interest: the estimation for the amplitude of the effect itself (e.g., a *β* value from regression analysis or GLM) and the associated statistic (e.g., *t* or *z*). The former provides the magnitude of a physical measurement, which is the essence of scientific investigation, while the latter offers statistical substantiation for the effect estimate in the form of a significance level (or confidence interval, the implied range that may contain the effect estimate with a certain likelihood). While the relationship between the two quantitates is tight, each conveys distinct information about the result of the experiment; in most scientific disciplines, it is considered unacceptable if only significance is reported (Sullivan and Feinn, 2012): the statistic value serves as auxiliary evidence for the existence of the targeted effect, and it is the effect estimate itself that is the center of investigation as the physical property of interest. For example, suppose that physicists would like to validate the predictions of the general relativity (Einstein, 1915) by investigating the gravitational waves from the merger of two black holes. It would be hard to imagine that they would only report a statistical value or the significance of their measurement (e.g., a chance of 1 event per 203,000 years, or a significance level of 3.4 × 10^−7^), but that they would not reveal the strength of the signal they have detected (a peak gravitational-wave strain of 1.0 × 10^−21^ in the frequency range of 35 to 250 Hz) (Abbott et al., 2016).

However, within the field of neuroimaging, it has remained the predominantly common practice to report only statistical mapping tests in publications and presentations, a custom which has been largely (and perplexingly) immune to critical scrutiny. For instance, one typically sees brain results provided as blobs whose color spectrum corresponds to *t*- or *z*-values (or occasionally to *p*-values), and most of the time the underlying degrees of freedom are left out, rendering the statistics even harder to interpret. Similarly, in tabulated results for brain regions, standard reports usually contain the coordinates and statistic value at a single peak voxel (which is itself defined, again, as the maximum of the statistical values, not of the effect estimates, within the region), and the effect estimate at such a peak voxel is rarely reported. The same phenomenon commonly occurs in reporting results of seed-based correlation analyses for resting-state data, where the brain maps and tables usually show the statistic (often *z*) values instead of and without including inter-regional correlations.

Recently there have been a number of discussions about the use and misuse of *p*-values in the scientific community (e.g., Wasserstein and Lazar, 2016; Nuzzo, 2014), and others have been more critical of the “cult” or “obsession” of statistical significance (e.g., Ziliak and McCloskey, 2009). The editors of the journal, Basic and Applied Social Psychology, have gone so far as to take the seemingly extreme step as to no longer accept papers with *p*-values due to the concern of the statistics being used to support lower-quality research (Trafimow, 2014). In a sense, our concern here is related, and addressing it would also alleviate many of these other topical issues, but the issue is specifically focused on the need for including the effect estimate in neuroimaging studies. To frame the discussion here, we quote the six guiding principles on *p*-values in a recent statement released by The American Statistical Association (ASA) (Wasserstein and Lazar, 2016):

1. *P*-values can indicate how incompatible the data are with a specified statistical model.
2. *P*-values do not measure the probability that the studied hypothesis is true, or the probability that the data were produced by random chance alone.
3. Scientific conclusions and business or policy decisions should not be based only on whether a *p*-value passes a specific threshold.
4. Proper inference requires full reporting and transparency.
5. A *p*-value, or statistical significance, does not measure the size of an effect or the importance of a result.
6. By itself, a *p*-value does not provide a good measure of evidence regarding a model or hypothesis.

We believe that the neuroimaging field needs to move forward to promote the reportage of the effect estimates along with the corresponding statistics. We first discuss the statistical terms in the context of FMRI analyses, highlighting specific features related to that field. We then argue that full reporting in FMRI is necessary and promotes good scientific practice, clarity, increased reproducibility, cross-study comparability and allows for proper meta and power analyses. Finally, we provide several recommendations for researchers and software designers to facilitate these “best practices” actions.

## What is the effect estimate in neuroimaging?

In neuroimaging, the ultimate focus is on the physical evidence for the brain’s neuronal response, which evidence is typically embodied in the strength of the FMRI BOLD signal. For task-related experiments, the response strength is reflected in the effect estimate (or *β* value) associated with a task/condition or with a linear combination of *β*’s from multiple tasks, such as the contrast between two tasks. For seed-based correlation analyses with resting-state data, time series correlation captures the relationship between a seed and the rest of the brain. Similarly, for naturalistic scanning, one measure is the “inter-subject correlation” (ISC) at a region that features the synchronization or similarity among subjects (Hasson et al., 2004). Here, we use the term “effect estimate” to refer generally to any of these or analogous cases: the estimated response magnitude (e.g., *β* value) of a regression model or GLM, the estimated correlation coefficient in the context of correlation analyses, etc.

We note that in the statistical literature, the phrase “effect size” can typically encompass two distinct scenarios: one for describing absolute effect size (the estimated magnitude of an effect under investigation, e.g., sample mean or the estimated *β* in a regression model), and the other for describing standardized effect magnitude (e.g., Cohen’s *d*), which is typically used when the measurement units have no intrinsic meaning (e.g., Likert-type scale adopted in survey research), when a comparison is performed between two different scales (e.g., relative effect sizes among different confounders such as age and sex), or when data variability is the focus of study (Sullivan and Feinn, 2012). While it is well known that the acquired BOLD signal has only arbitrary units, therefore it might seem that the second usage of effect size is a good candidate. However, FMRI data are commonly scaled to a more meaningful evaluation in terms of percent signal change (as discussed further below). As such, here we use the term “effect estimate” in FMRI to refer to the unit-bearing case of “effect sizes” in the context of percent signal change.

## What does a *t*-statistic value reveal in neuroimaging?

A *t*-statistic value for an effect estimate is calculated as the latter divided by its standard error, which represents the reliability or accuracy of the effect estimate. Thus, the *t*-statistic is a mixture of the effect estimate and the noise estimate, and there is little reason to think that the noise estimate is directly relevant to neuroscience. As a dimensionless measure, the *t*-statistic is more susceptible to sample size (number of trials or subjects), signal-to-noise ratio (SNR), preprocessing steps/methods, experimental designs, unexplained confounds, and scanner parameters than the effect estimate itself. Therefore, statistic values only serve the purpose of a binary inference of null (e.g., there is no difference between the two conditions) versus alternative (e.g., there is difference between the two conditions) hypotheses, and it does not provide any information about the specific response magnitude. For example, two voxels (or regions) with the same *t*-statistic value in the brain do not mean the same response amplitude, and *vice versa* (Fig. 1). That is to say, the *t*-statistic does not carry enough interpretation information for the effect of interest.

**Figure 1:**
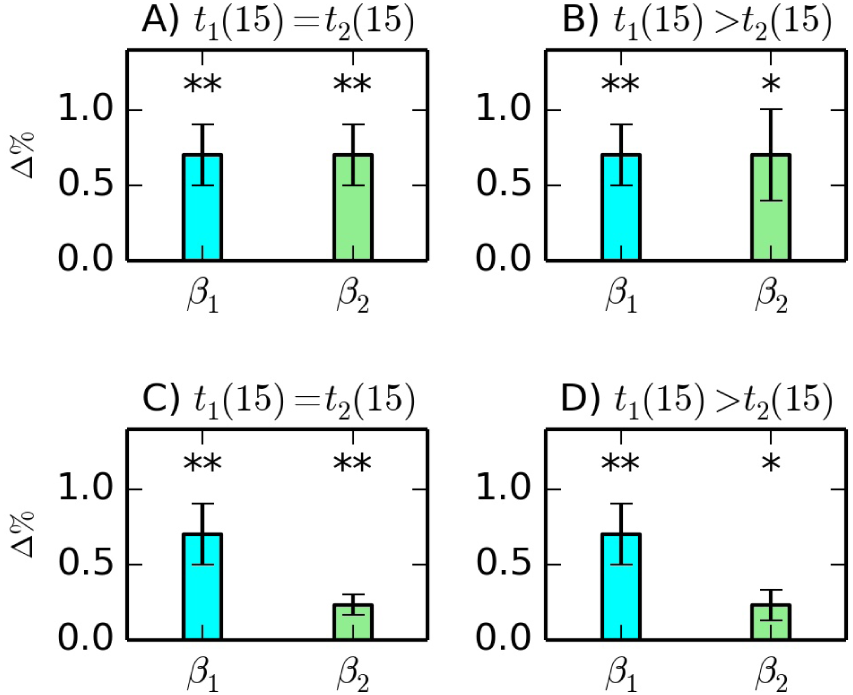
A statistic value alone does not reveal the relative magnitude for an effect of interest. Specifically, two identical *t*-values (here, with 15 degrees of freedom) may have similar (A) or dramatically different (C) effect estimates. On the other hand, two different *t*-statistic values may have the same (or opposite) sequence as (or to) that of the corresponding effect estimates; for instance, a larger *t*-value could correspond to a larger effect estimate if the standard error is roughly proportional to the effect estimate (D) or similar or even smaller effect estimate if the standard error is smaller (B). The numbers inside the parentheses are the degrees of freedom for the *t*-statistic, and asterisks indicate orders of magnitude in *p*-values: * 0.01 ≤ *p* < 0.05; ** *p* < 0.01. Effects are scaled units of percent signal change.

## Practical realities/difficulties of FMRI

There are several features inherent to FMRI acquisition and analysis that present challenges to an investigator interpreting and reporting results. At first glance, some of these may seem to explain the present practices of reporting only statistic values as results. We describe them briefly here, and then discuss how they actually necessitate, rather than discourage, the inclusion of effect estimates in the end.

### Units and scaling

As noted above, one complication of the FMRI signal is that the numerical value from the scanner does not have any specific physical meaning and is essentially arbitrary. As a consequence, the signal value may vary across brain regions, sessions, days, subjects, studies, and scanners. To deal with this arbitrariness, a normalization step is typically adopted by researchers by scaling the signal so that the relative magnitude of the BOLD response is comparable between different contexts. For example, by default in AFNI (Cox, 1996) the time series is scaled by the mean value at each voxel, so that the effect estimate can be directly interpreted as a percent signal change relative to the voxel-wise temporal mean; as a result, effect estimates themselves are interpretable, carry real information about the size of the BOLD effect, and are comparable across brain regions, conditions, subjects, groups, studies and scanners^1^.

One may argue that the voxel-wise baseline, instead of the mean, is a more accurate candidate to serve as the scaling factor. However, in FMRI the drift effect (or the presence of low frequency components due to scanner drift, shim effects) embedded in the signal complicates the isolation of the “real” baseline value. In practice, the fluctuations due to the task effect are very small relative to the absolute values of the signal (e.g., most task effects are around 1% or less relative to the BOLD signal mean), leading to a negligible difference when the voxel-wise mean, instead of the “true” but unknown baseline, is used in scaling^2^. Even if there are different preferred mechanisms of scaling, it appears to be a truth universally acknowledged that the BOLD signal can and should be calibrated through a normalization step, providing a meaningful and comparable measure. While there is not a single method for calibrating the effect estimate or signal change to a meaningful unit that is uniformly adopted by all researchers, such a difficulty should not be an excuse for not reporting the BOLD response.

### Modeling difficulties

One aspect of FMRI data is that the hemodynamic response (HDR) is captured by a curve with a slow upstroke and a sluggish recovery; the curve may also contain an undershoot right after the stimulus onset or at the end of the recovery phase (D’Esposito et al., 1999). In addition to the overall amplitude, the response may vary across cognitive states, tasks, brain regions, and subjects with respect to response characteristics such as rise and fall speed, peak duration, undershoot shape, and overall duration. The nature of the HDR is still not fully understood due to the complicated and multifaceted biophysical processes involved.

As the underlying components comprising the BOLD signal are still poorly understood, the performance of the regression model at the individual subject level is often poor. For example, attenuations across trials or within each block are usually not considered; the impact of physiological (cardiac and breathing) effects is mostly lacking, though it is occasionally modeled (e.g., ANATICOR, Jo et al., 2010). Because of these factors, the variance due to poor modeling overwhelms all other sources (e.g., across trials, runs, and sessions) in the total data variances (Gonzalez-Castillo et al., 2016); that is, the majority (e.g., 60-80%) of the total variance in the data is not properly accounted for in statistical models. There are also strong indications that a large portion of BOLD activations are usually unidentified at the individual subject level due to the lack of power (Gonzalez-Castillo et al., 2012). The detection failure (false negative rate) at the group level would probably be equally high, if not higher. Due to the presence of large variability and unaccounted-for noise, low reliability leads to inaccurate estimation of the effect of interest.

Another modeling difficulty that arises when comparing effect estimates across studies is the dependence of the BOLD effect percent signal change on scanning parameters (e.g., B_0_, TE, slice thickness, etc.). The current state of modeling does not make combining/contrasting effect estimates from significantly different types of scans practicable. For this reason, it is important to clearly specify the MRI setup used.

## Limitations of statistical significance testing

Under the methodology of null hypothesis significance testing (NHST), the statistic value is mainly used to determine the statistical significance level of an effect estimate so that false positive rate is controlled. Once the value surpasses the threshold, the specific value of the statistic is neither as informative nor as important as the response amplitude or effect estimate. The current misplaced focus on statistical significance when reporting a scientific result (Ziliak and McCloskey, 2009) is equally detrimental as shown by a popular statistical fallacy: If the result is not statistically significant, then it proves that no effect or difference exists. As the *p*-value under a null hypothesis is a conditional probability, it cannot be stated that the probability of obtaining the data under the current study given the null is the same as that of the null given the data.

There is a clear difference between statistical significance and practical significance. The absence (or ignorance) of a real effect estimate in results reporting has prompted the distinction between the two types of significance: substantive significance or practical significance in terms of effect magnitude and statistical significance in terms of probability threshold (Gelmen and Stern, 2006). For example, it was shown that “emotional contagion occurs without direct interaction between people (exposure to a friend expressing an emotion is sufficient), and in the complete absence of nonverbal cues” through Facebook (Kramer et al., 2013). However, it was later pointed out that the effect size measured by Cohen’s *d* = 0.02 was so small that such a tiny difference in emotional contagion is not practically meaningful. In other words, a trivial effect (a tiny difference between two groups or conditions, or a negligible correlation) can become statistically significant with enough sample size. For example, a drug effect in a clinical trial, even if statistically significant, may not offer much practical benefit when the effect is small (e.g., lowering cholesterol level by 2.7 mmol/L). Similar pitfalls have been seen in studies which “demonstrated” that beautiful parents have more daughters, and violent men have more sons (Gelman and Weakliem, 2009). Importantly, without presenting the effect estimate, not only would one be unable to gauge the false negative rate or power of the study, (i.e., the probability of failure or success, respectively, to detect the effect), but it would also be impossible to assess two other useful but less known errors (Gelman and Tuerlinckx, 2000): type M (tendency to over- or under-estimate the effect magnitude) and type S (likelihood of obtaining the incorrect directionality or sign of the effect).

Activation identification in FMRI data analysis heavily relies on contrasting between conditions; however, another subtlety is that the contrast between a significant effect and a nonsignificant one is not necessarily itself statistically significant. For example, suppose that, with 16 subjects (and 15 degrees of freedom), positive and negative conditions have effect estimates of 1.0 and 0.45 percent signal change, respectively, and both estimates have the same standard error of 0.3. Even though the positive condition is statistically significant (*t*(15) = 3.33, two-tailed *p* = 0.0045) and the negative condition is not (*t*(15) = 1.5, two-tailed *p* = 0.15) at 0.05 level, their contrast could be statistically insignificant (e.g., *t*(15) = 1.65, two-tailed *p* = 0.12) (Fig. 2).

**Figure 2:**
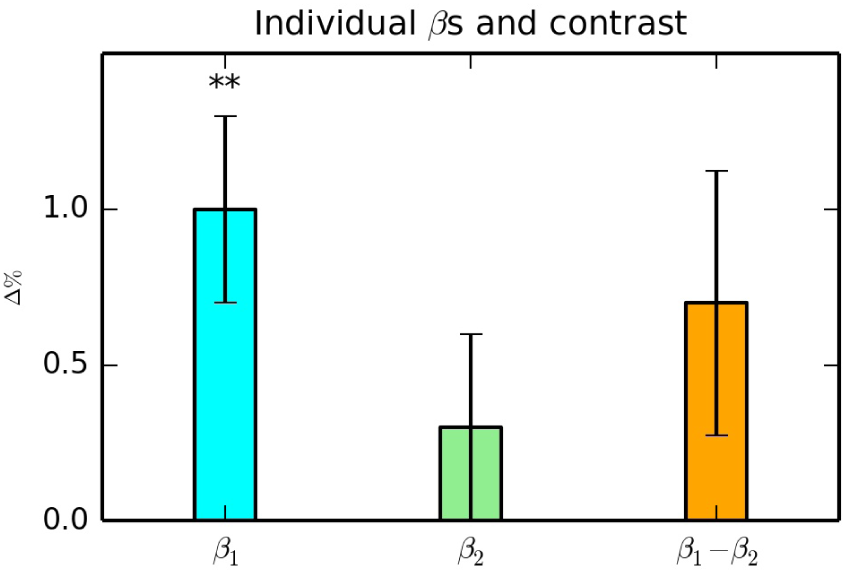
A statistically significant (blue) and insignificant (green) effect are shown both in scaled units of percent signal change. However, their difference might be practically significant but not statistically significant (yellow). Asterisks indicate orders of magnitude in *p*-values: * 0.01 < *p* < 0.05; ** *p* < 0.01.

The classical statistical testing is consistent with the Popperian paradigm in which science advances through the proposition and refutation of hypotheses (Popper, 1963). However, the omnipresence of focus on statistic values alone, while ignoring the effect estimates, unavoidably encourages and facilitates a yes/no binary thinking, and has in fact led to the false interpretation that sub-threshold regions have no activation and that supra-threshold regions comprise the entire story (Gelman, 2013). In addition, the approach suffers from a “statistical significance filter” (Gelman and Weakliem, 2009): results that reach a preset significance level inherently overestimate the effect and also tend to go in the wrong direction.

## Why is it crucial to report effect estimates?

The effect estimate provides a piece of hard, quantitative evidence in an analysis, and it should be reported as the main finding of a modeled or measured effect (Sullivan and Feinn, 2012). The corresponding statistic or *p* value usually indicates the reliability or accuracy of the effect estimate, but it cannot replace the information content of the effect estimate itself. For this reason, the importance of reporting the specific effect estimate under study has been repeatedly emphasized in various fields. For example, one recommendation from the American Psychiatric Association (Wilkinson et al., 1999) reads: “Always present effect sizes for primary outcomes... If the units of measurement are meaningful on a practical level (e.g., number of cigarettes smoked per day), then we usually prefer an unstandardized measure (regression coefficient or mean difference) to a standardized measure (*r* or *d*).” We enumerate here specific examples and applications of this principle within the FMRI context.

### Reproducibility

Reproducibility is critical for scientific investigations, and it can be quite challenging for FMRI studies, as the data typically have low SNR and low reliability for each effect estimate. One should not overemphasize the statistical thresholding and lose sight of the scientific context, particularly where the noise is usually much stronger than the signal in the data. In recent surveys, about 60% of published experiments failed to survive replication in psychology (Baker, 2015) and about 40% in economics (Bohannon, 2016), and the situation with neuroimaging is likely not much better (Griffanti et al., 2016).

In fact, the availability of the effect estimate in the literature becomes pivotal in cross-examining or reproducing the results across studies. Verification for regional activations based on statistical significance would partially serve the purpose, but reproducibility cannot be solely built on statistical values. The notion that statistical significance alone does not imply result replicability is nicely captured by Thompson (1999): “it would be the abject height of irony if, out of devotion to replication, we continued to worship at the tabernacle of statistical significance testing, and at the same time we declined to (a) formulate our hypotheses by explicit consultation of the effect sizes reported in previous studies and (b) explicitly interpret our obtained effect sizes in relation to those reported in related previous inquiries.”

With both the effect estimate and its standard error (or reliability, which is embedded in the *t*-statistic value, for example) available, one can readily compare the effect estimates across conditions, regions, subjects, groups, studies, scanners, etc. For example, suppose that a previous study indicated an effect estimate of 0.73% signal change with a statistic value of *t*(16) = 4.12 at a peak voxel (defined by the maximum effect estimate within a cluster). In such a case, a researcher would find that having an effect estimate of 0.65% with *t*(22) = 3.75 in her own study would be compatible with the existing result, while an effect estimate of 0.1% with *t*(22) = 3.35 would unlikley be. Obviously such comparisons (or reproducibility) would be impossible if only statistic values are reported in the literature, as currently prevalent in neuroimaging.

Furthermore, one can also use effect estimate reporting to easily spot unrealistic results at a region, either in one’s own pre-published work or, an unfortunate practical necessity, in an existing research article. For example, a region might show up having more than 3% signal change while still exhibiting a reasonable statistical significance due to modeling issues, noise, etc. If only statistics were used for thresholding, coloring and reporting, then such an artifactual result would likely go undetected by either the authors or, later, other readers. Thus, viewing the effect estimates themselves provides an extra layer of safety against false positives, increasing reproducibility in reporting.

### Clarity

It is a common practice in FMRI literature to present brain activation maps that are both thresholded and colored by statistic values. However, such presentations entirely ignore the effect estimates, and such coloration has been shown to lead to distorted impression of the results in recent surveys (Engel and Burton, 2013). If only the significance level of a correlation or BOLD response at a region is given, one would have no idea about the strength of the effect or the association, and thus the scientific relevance is missing. In other words, with the current practice of reporting statistic values alone, at best the results are ambiguous and at worst they are misleading.

To drive home the point that a statistic or p value is not the whole picture nor as informative as combining with the effect estimate, consider the following example. Suppose that at one region the effect estimate is 0.03% signal change with p = 0.001 while at another region the response is 0.94% with *p* = 0.053. Is the higher statistical significance with the first voxel more worthy of reporting than the second? On the surface, the response of 0.03% at the first region occurred with greater confidence while the second region failed to reach the arbitrarily designated significance level of 0.05. However, the response magnitude of 0.94% is quite a bit stronger and might be more neurologically relevant or important than the statistically significant response of 0.03%. Furthermore, the second region might have reached the nominal significance level with a larger number of subjects. Looking at this example without the effect estimates, one might easily misinterpret the results.

Directly relevant to the neuroimaging community is the moral from these examples: without the effect estimate, the sole focus on statistical significance often presents a distorted picture. Specifically, the power with neuroimaging data is typically low due to the the facts that large parts of the signal that cannot currently be accounted for and that there is large variability across subjects. The presence of many false negatives may lead to the illusion that a statistically insignificant effect is equivalent to a nonexistent effect, when in some cases there are not enough data to discern whether the effect is practically important. In other words, type M errors tend to increase, and a distorted interpretation may occur without the presence of effect estimates that may be assessed more accurately than the decontextualized statistic values.

### Validation of BOLD response detection power through effect estimates

Although most research-oriented investigations place a heavily-lopsided emphasis on the false positive rate controllability, sensitivity (or power) may also be a primary focus under some circumstances, such as pre-surgical detection, where the efficiency is usually less than 10% (Button et al., 2013). Several particular factors may contribute to a cluster not being able to achieve the desired significance at the group level under a rigorous procedure.

a. To achieve the desired significance or power at the cluster level (or in the FDR sense), it is usually necessary to have a large number of subjects, which most studies lack due to financial and/or time costs.
b. Spatial alignment has multiple steps including cross-TR (“motion correction”), cross-session, cross-modality and cross-subject components, increasing the overall chance of misalignment. An erroneous or even suboptimal alignment procedure will surely impact the power performance at the group level.
c. The variation in response magnitude or SNR across regions, as well as the variation of the underlying region’s spatial extent, may also lead to different efficiency in activation detection across the brain. An intrinsically small response magnitude or small region, such as the amygdala, requires a smaller voxel-wise *p*-values to survive the family-wise error (FWE) or false discovery rate (FDR) correction compared to their larger counterparts, and this may not always be realistic to achieve in a study. The popular small volume correction (SVC) is offered as a band-aid solution, but is not always rigorous or valid, and may become problematic when other regions are of interest at the same time.
d. If a two-tailed test, when appropriate, is strictly performed instead of two separate one-tailed tests as typically practiced in the field^3^, or if FWE/FDR correction is rigorously executed, many studies would rightly face the issue of power deficiency.

The issue of reporting marginally significant effects is controversial (e.g., Pritschet et al., 2016). Should one not report a cluster simply because it cannot pass the rigorous statistical thresholding through FWE/FDR control at the present group size? We argue that, even if a cluster fails to survive rigorous correction, it does not necessarily mean that the results are not worth reporting, because they may be suggestive and provide some benchmark for future confirmation. Statistical inference should not be a binary decision, and the inclusion of effect estimates allows for a consistent approach to avoid this and to achieve a balance between false positives and false negatives (Lieberman and Cunningham, 2009). Thus we propose a two-tier approach to reporting clusters. In addition to the conventional FWE control, we believe that, if the individual voxels within a region achieve a basic significance level (e.g., *p* ≤ 0.05) and if the cluster possesses some practically significant spatial extent, its reporting is warranted. Nevertheless, the reporting has to be combined with the corresponding effect estimate as well as a cautionary statement about the marginality. On the other hand, the activation of a cluster may become questionable with an unreasonable effect magnitude (e.g., 3.5% signal change) even if the cluster survives stringent statistical thresholding, and again, readers can only detect such suspicious results if the effect estimate is reported, providing a safeguard against potential false positives (Fig. 3).

**Figure 3:**
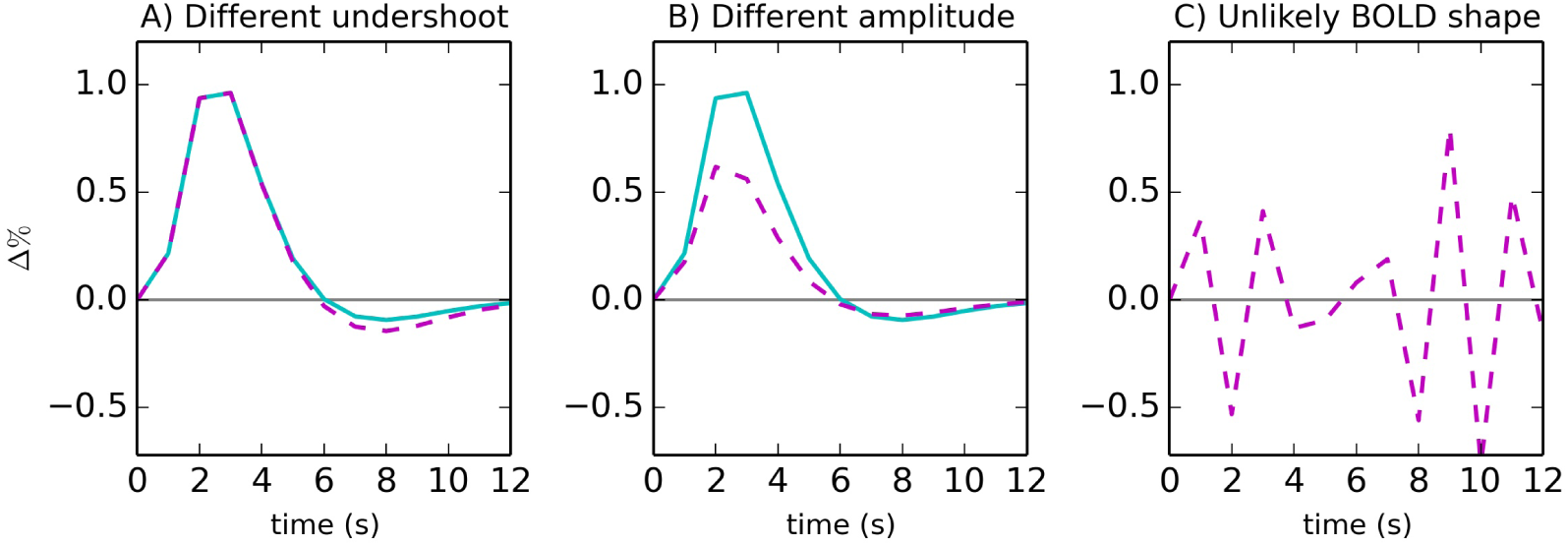
Modeling with multiple basis functions may provide more accurate characterization of the HDR as well as more powerful activation detection. For example, differences in shape features such as undershoot (A) and peak/recovery duration can be readily revealed in addition to peak (B). Furthermore, a false response curve, although statistically significant, would be identified (C) if its estimated shape dramatically differs from the signature shape of HDR.

### Validation of BOLD response modeling through hemodynamic response curve

There are three common approaches to modeling the BOLD HDR. The first one presumes a fixed shape (or model-based) impulse response (IRF), such as the gamma variate in AFNI (Cohen, 1997) or the “canonical” IRF in SPM and FSL (Friston et al., 1998a). With this method, a single regression coefficient (or β) associated with each condition in the individual subject analysis reflects the major HDR magnitude (e.g., percent signal change). The second approach makes no assumption about the IRF’s shape and estimates it with a set of basis functions, the number of which varies depending on the basis set and the duration over which the response is being modeled. For example, a common approach to this estimated-shape method consists of using a set of equally-spaced TENT (piecewise linear) functions (linear splines), and each of the resulting regression coefficients represents an estimate of the response amplitude at some time after stimulus onset. This produces an ordered set of effect estimates for each modeled HDR. The third approach lies between the two extremes and uses a set of two or three basis functions (Friston et al., 1998b). In this adjusted-shape method, the first basis (canonical IRF) captures the major HDR shape, and the second basis (the time derivative of the canonical IRF) provides some flexibility in modeling the delay or time-to-peak. The third basis (resulting curve, which is the derivative relative to the dispersion parameter in the canonical IRF) allows the peak duration to vary. Here, as well, multiple effect estimates are associated with a single HDR.

With only a single parameter per condition, the fixed-shape approach is the most efficient and statistically powerful among the three, if the presumed shape is reasonably close to the ground truth. This technique is widely adopted because the corresponding group analysis is the easiest. With the adjusted-shape method, the common practice at the group level is to focus only on the first effect estimate, ignoring the shape information captured by the second and third coefficients. Group analysis using multiple basis functions has recently been extensively explored (Chen et al., 2015), and the HDR shape information in the sequence of effect estimates can be carried from the individual level over to the group level. The powerful validation aspect of this approach is that, even if a region is marginally significant, the investigator may argue for the existence of an effect with the presence of the signature shape of HDR curve, as well as for subtle response differences in the undershoot, recovery phase, etc. The graphical representation of HDR profiles (see Fig. 3) gives one a reassuring observation or an extra confidence about their reliability that could not be gained only through the conventional statistical safeguards (e.g., when a cluster fails to pass rigorous thresholding). With the availability of effect estimates at the multiple time points of the whole HDR, it would be hard to fully deny the suggestive value of reporting the cluster together with its effect sizes and HDR profiles.

### Meta analysis and power analysis

As an integration approach, meta analysis in FMRI is usually performed to combine and summarize the results from various studies that are importantly not necessarily fully consistent with each other. There have been multiple methods developed for meta analysis. For example, the summarization may be based on voxel-wise results, a specific region (ROI), labels, coordinates, image, or activation likelihood estimation (Radua and Mataix-Cols, 2012). Most of the existing methods do not consider the effect estimates, in large part because such information is missing in the literature.

FMRI studies incorporate many factors that easily vary across sites, such as sample size (e.g., number of subjects and number of repetitions for each condition), specific task designs, scanners, etc.; and, as a result, both the magnitude of an effect and its reliability could be largely heterogeneous across reports. If the synthesis through meta analysis is solely based on coordinates or statistic value, the results could be unreliable. A recent study has shown that, when both effect estimates and their standard errors (which can be derived from the *t*-statistics) are available, meta analysis through a mixed- or random-effects model (Maumet and Nichols, 2016) would be more robust than other alternatives such as label- and coordinate-based approaches (e.g., coordinates only: activation likelihood estimation, Eickhoff et al. 2012; coordinates and Gaussianized *Z*-values: Radua and Mataix-Cols, 2009; Costafreda et al., 2009; Yarkoni et al., 2011). Furthermore, if those studies in which a region marginally survives (or even fails to survive) the FWE correction at the cluster level are included, an approach with both effect estimates and their stability information incorporated in the meta analysis would be more immune to publication bias.

The effect estimate is also a necessary quantity for power analysis. To design an experiment, the investigator may take information from previous studies and use power analysis to either 1) determine the sample size required to achieve a preset power (or false negative rate), or 2) assess the power of a given study (how likely one would detect a specific effect magnitude under a particular context). For both calculations, the statistic value as well as the effect estimate are needed as prior information. Even though mostly power analysis is currently performed with the peak value of *t*-statistic in the brain or a region (Durnez et al., 2016), the approach can be improved if the effect estimates are available in addition to statistic values. For example, the peak defined by the effect estimates within a cluster instead would be a more accurate representation than one by the *t*-statistic values. In addition, the availability of effect estimates would allow the investigator to perform conventional power analysis at the voxel, instead of region, level.

Looking forward, as the amount of public data and subsequent cross validations, meta and power analyses increases, it is vital to start providing results from more robust results for agglomerative approaches.

## Recommendations and conclusion

Scientific investigations usually involve data collection from observational studies or meticulously-designed experiments. Raw data with no or little extraction and compression would clutter or even obscure the intended message from the investigator. On the other hand, overly summarized data or missing information would present less convincing conclusions, or, worse, lead to misleading impressions. Statistic values alone do not represent the whole scientific endeavor, and there is no reason to believe that neuroimaging should be an exception in which physical measurement is largely ignored. As a crucial part of scientific investigation, good statistical practice should reveal relevant quantitative components of data summarization including the amplitude of brain response in neuroimaging. Such numerical and graphical information would promote reproducibility and aid power and meta analysis. In addition, the effect estimate may either offer extra support to or counter the interpretation made from the statistical significance alone; either case leads to more accuracy, and therefore its inclusion should be reassuring to researchers.

As an antidote to *p*-hacking or the obsession with statistic values, complete rejection of *p*-values in scientific reporting would likely be an overreaction. We believe that it would be equally impropriate to report only the effect estimate without the auxiliary information about its reliability in the form of standard error, confidence interval, or statistic value. Both pieces of information are needed to see the whole picture. In addition to the response magnitude’s serving as a benchmark, another benefit is that, if these multiple pieces of information were available in literature, one could identify those regions that showed substantial response magnitude but failed to achieve a significance level in the study due to large variability across subjects (such results are typically undisclosed.

Some effort has been devoted to promote the standardization of the reporting process in neuroimaging analysis (e.g., Poldrack et al., 2008; Carp, 2012; Nichols et al., 2016), though the important issue of reporting effect estimates has not been paid much attention. In this commentary, we have argued that reporting effect estimates has the same goal and benefit as standardization and that it is in fact necessary in order to improve results reporting in the field. In addition to revealing modeling specifics such as all explanatory variables, the number and directionality of post hoc tests, we strongly believe that effect estimates (e.g., in a scaled unit such as percent signal change) should be reported along with statistic values, instead of having excessive focus only on the latter in graphical representation. In addition, reporting the standardized effect (e.g., Cohen’s *d*) may be a valid alternative as well.

Regarding clusterization, we recommend that:

1. the statistic values be used for thresholding only (not for colorization, determining maxima of activity, etc.);
2. the activation patterns in brain images be colored by effect estimate values (e.g., percent signal change, correlation), not by statistic values; and
3. the full set of parameters (threshold value, degrees of freedom for each statistic test, cluster-wise probability, etc.) be explicitly stated.

Effect estimates should also be included in tabulated results at the regional level, with the peak defined as the maximum of the effect estimate, not of the statistic values. They can serve as another layer of supporting evidence in activation identification, and this becomes especially crucial when some practical constraints (e.g., few subjects, suboptimal spatial cross-modality/subject alignment, small regions) lead to a situation in which a cluster fails to survive rigorous thresholding. Analytical toolboxes and software should facilitate, nurture, or even enforce a standardized process of generating proper and complete results reporting, thereby reducing the emphasis of *p*-values.

Our suggestions are aligned with and complementary to a proposal of avoiding misinterpretations through graphical representation of confidence intervals (Engel and Burton, 2013), as well as the guiding principles regarding reporting statistics in the recent ASA statement (see Introduction; Wasserstein and Lazar, 2016). Einstein noted that, “It can scarcely be denied that the supreme goal of all theory is to make the irreducible basic elements as simple and as few as possible without having to surrender the adequate representation of a single datum of experience” (Calaprice, 2010). Within the applied field of FMRI, this notion of making results “as simple as possible *but not simpler*” should be taken to heart and adopted as well. We feel that this can be done only by including the full model reports of effect estimates and statistics in the literature.

## Footnotes

1 Similarly, “grand mean scaling” is typically performed in FSL (http://fsl.fmrib.ox.ac.uk/fsl/fslwiki/) and SPM (http://www.fil.ion.ucl.ac.uk/spm/), by dividing the signal by the average value across the brain as well as across time. The purpose of grand mean scaling is to bring the effect estimates to a similar range so that they are roughly comparable across brain regions, sessions, days, subjects, studies, and scanners. However, such a scaling method does not exactly lead to the interpretation of percent signal change because of spatial heterogeneity. A separate toolbox MarsBaR (Brett et al., 2002) is often used to convert the effect estimates into percentage at the regional level.

2 The negligible effect of replacing the true “baseline” value by the voxel-wise mean can be demonstrated by a back-of-the-envelope calculation. Suppose that the signal intensity at a voxel has a mean value of 2400 for the time series (after slow drift effects are removed), peak intensity corresponding to a task is 2410, and a “real baseline” value is 2390. The scaled peak value at the voxel by the mean is 100 × 2410/2400 ≈ 100.417, and the scaled baseline value of 100 × 2390/2400 = 99.583. The percent signal change for the task relative to the baseline is thus estimated as (100.417 − 99.583)/100 ≈ 0.834% in the regression model. Alternatively, if we analyze the data without scaling, the “true” percent signal change of the condition would be calculated as (2410 − 2390)/2390 ≈ 0.837%. The ratio of the difference between the two estimates relative to the true effect estimate is (0.837 − 0.834)/0.837 ≈ 0.358%.

3 When a two-tailed *t*-test is more suitable, blindly performing a one-tailed *t*-test is equivalent to artificially inflate the *p*-value by a factor of 2; for example, a *p*-value of 0.01 for a two-tailed *t*-test corresponds to a *p*-value of 0.005 for each of the two associated one-tailed *t*-tests.

## Acknowledgments

The research and writing of the paper were supported by the NIMH and NINDS Intramural Research Programs (ZICMH002888) of the NIH/HHS, USA.

